# A robust, sensitive phylogenetic method enables gene-level metagenomic analyses

**DOI:** 10.64898/2026.07.15.738679

**Authors:** Nghi Tran, Kathryn Kananen, Patrick H. Bradley

## Abstract

A key goal in the microbiome field is to move from taxonomic associations towards mechanistic hypotheses about microbial gene function. However, most methods for linking microbiome changes to specific genes are biased towards finding marker genes, with weak evidence for functional relevance. Phylogenetic regression can address this issue and has been previously applied to changes in microbial prevalence, but many environments (such as the gut in health vs. disease) are characterized more by changes in abundance, which presents unique statistical challenges. We show that when applied to real differential abundances from metagenomes, phylogenetic regression has an anti-conservative bias, indicating inflated false positives. We develop an alternative non-parametric method called “robust permutration,” designed specifically for differential abundance data, and evaluate its performance against phylogenetic regression as well as several other phylogenetic comparative methods in realistic simulations of metagenomic data. These results show that robust permutration is the most powerful method that appropriately controls the false positive rate. We further apply robust permutration to a human case-control study of liver cirrhosis, revealing that *Lachnospiraceae* abundance in disease is linked to a previously uncharacterized iron-sulfur transcription factor encoded near homologs of the butyryl-CoA oxygen oxidoreductase system, a recently discovered system for oxygen detoxification. This illustrates how robust, sensitive phylogenetic methods can enable the generation of new molecular hypotheses directly from metagenomic case-control data.

**Importance:** Previously, we showed that phylogenetic regression can effectively detect genes associated with microbial presence or absence while correcting for evolutionary relationships. Unexpectedly, however, we here observe that this method can lead to high false positive rates when applied to microbial abundance data. In realistic simulations, other methods we test either have similar problems with false positives, or display very low power. We outline a new statistical test that better accounts for measurement uncertainty, outliers, and model violations, achieving more balanced sensitivity and accuracy than competing methods. Applying this test to a cirrhosis study reveals an uncharacterized transcription factor enriched in disease, with an apparent role in oxidative stress based on its sequence and gene neighborhood. This suggests a functional explanation for the observed taxonomic shifts, and demonstrates how improved phylogenetic methods could help inform future microbiome-targeted treatments.

## Introduction

High-throughput DNA sequencing has transformed our ability to measure changes in microbial communities. In host-associated microbiomes, these changes have been linked to a wide variety of health outcomes, including human liver disease^1,2^, metabolic syndrome^3^, and autoimmune disorders^4^. Many of these host phenotypes can be altered by antibiotic treatment or transferred via microbiome transplants, pointing towards a causal relationship. In free-living environments, microbial communities shape the biosphere through their metabolism^5^, which helps drive key nutrient cycles^6^ in ways that can affect outcomes like the production of greenhouse gases^7–9^. However, our ability to intervene in the microbiome is still largely limited to global perturbations, such as transplanting communities or treating with antimicrobials, which have a high potential for unintended consequences. Antibiotic treatment in hosts has been linked to increased risk for pathogen colonization^10^ as well as conditions like metabolic syndrome^11^; in the environment, antimicrobial contamination is linked to the spread of resistance^12^. In hosts, microbiome transplants require healthy donors, posing problems for scaling the treatment, especially as donors must be screened rigorously to avoid potentially deadly sequelae^13^. There are also recent indications that transplants can lead to persistent anaerobic colonization of the small bowel^14^. It is therefore a high priority across the microbiome sciences to move from taxonomic associations to a molecular understanding of what drives the changes we observe.

How can we do this effectively across environments, including in cases where we may have weak prior knowledge about microbial gene function? One way is to look for microbial genes that are statistically associated with biome-level phenotypes, an approach that has analogies to (for example) human genetics. This has the advantage of not requiring cultivation or experimental genetic tools. Furthermore, because statistical associations can be untargeted, they can discover links to genes that currently lack a known, annotated function^15^, which are common in the microbiome^16,17^. Untargeted approaches can help prioritize which of these are most worth characterizing, countering the “streetlight effect”^18,19^ where processes that are already well-studied receive outsized attention.

However, statistical approaches can also be susceptible to confounding by factors that are correlated with genotypes or phenotypes of interest without being causally related, leading to false discoveries. One such factor is phylogenetic correlation, which arises because closely-related species tend to share both genes and traits; this is analogous to population structure in genome-wide association studies^20^. Previous work has shown that phylogenetic correlation has a major impact on the analysis of metagenomic data, and that non-phylogenetic methods tend to find markers for clades that change, rather than genes with strong evidence for a functional relationship^21–24^.

So far, however, only two methods that explicitly consider phylogeny, *phylogenize* (version 1)^21^ and Phylogenetic Organization of Metagenomic Signals (POMS)^22^ have been developed specifically for the analysis of species-level metagenomic data (another recent method, Anpan^25,26^, also uses a phylogenetic comparative approach, but focuses on strain variation within a species). Methods specifically geared to metagenomic data are important as they present unusual statistical challenges, including mean-variance correlation^27^, compositionality^28^, and zero-inflation^29^.

The first version of *phylogenize*, which uses phylogenetic linear regression^30^, simplified this problem by only considering changes in microbial presence or absence. However, many changes observed in microbiomes, such as the human gut in disease, involve changes not just in presence but in abundance; dichotomizing the data throws away this information^31^. Furthermore, phylogenetic regression makes certain assumptions that may not be valid on real metagenomic data. One is that continuous traits evolve according to Brownian motion or a similar process; however, real traits may also have rapid shifts that are hard to model using this framework^32^. The presence of high-leverage outliers may also distort the regression fit^33^. POMS addresses some of these issues by using a novel non-parametric approach based on the phylogenetic isometric log-ratio transformation (PhILR^34^), allowing abundance information to be incorporated. However, simulations have suggested that POMS has lower sensitivity than phylogenetic regression^22^, particularly in cases where most important changes are recent (i.e., concentrated towards the tips of the tree). Additionally, it is not clear how POMS may be affected by factors like outliers or abundance data with unusual distributions.

Here, we show that standard phylogenetic regression can lead to biased results when applied to differential abundance data from real metagenomes, as well as to simulated data when outliers and model violations are added. Drawing inspiration from several different modern phylogenetic comparative approaches like ROBRT^33^, RRphylo^35^, and permulation^36^, we present an approach that combines robust phylogenetic regression with a new approach based on permuting evolutionary rates, which we call “robust permutration.” We apply this method with a tree transformation that allows us to model unequal measurement error across taxa. We show that robust permutration with per-taxon measurement error substantially reduces sensitivity to modeling assumptions and outliers, and that with an early stopping rule, it remains computationally tractable enough to test all gene families across multiple clades. To obtain reliable differential abundance estimates, we combine methods built specifically for microbiome data, like ANCOM-BC2^37^, with a general-purpose shrinkage approach (ashr^38^). In realistic metagenomic simulations, this new approach outperforms not only POMS but also other phylogenetic comparative methods that are not specific to the microbiome, like ROBRT^33^, phylogenetic regression with permulation^36^, and TreeWAS^39^.

Finally, we apply robust permutration to a case-control study of liver cirrhosis, focusing on the *Lachnospiraceae*, a highly prevalent but understudied clade whose abundance has previously been linked to health. The results show that *Lachnospiraceae* that are less depleted in cirrhosis are enriched for an uncharacterized iron-sulfur transcription factor from the Rrf2 family. This transcription factor tends to be encoded next to homologs of the recently described butyryl-CoA:oxygen oxidoreductase (BOOR) system for oxygen detoxification, as well as other anaerobic oxidative stress resistance genes like rubrerythrin and ribonucleotide reductase. This demonstrates that this method can be used productively to translate patterns of differential abundance among taxa in the microbiome into new molecular hypotheses.

## Results

### Standard phylogenetic comparative methods are not well calibrated on realistic microbial abundance data

The original implementation of *phylogenize*^21^ applied a phylogenetic linear model, implemented in the phylolm package^30^, to microbiome data. Briefly, *phylogenize* modeled the prevalence of taxa (or a measure of differential prevalence across environments, called “specificity”) as a function of gene content. In simulations under the null, that is, when there was no true gene-phenotype correlation, the phylogenetic approach controlled the false positive rate appropriately, while a standard linear model had false positive rates that could reach nearly 75%.

To make sure this method was appropriate for abundance data, we decided to perform a similar experiment using not only simulated phenotypes but also a “real” differential abundance phenotype. We calculated this phenotype by applying ANCOM-BC2^37^ to marine metagenomes from the Tara Oceans^40^ project (*Flavobacteriaceae* at the surface vs. the mesopelagic zone) and applying adaptive shrinkage^38^ (see Methods). Genes were simulated to have no true correlation with either phenotype (see Methods).

Surprisingly, while the phylogenetic linear model performed as expected for the Brownian motion (BM) simulation under the null model, showing a uniform distribution of p-values, it exhibited anti-conservative behavior for the real phenotype data, indicating a loss of false positive control (**Figure 1A**). This suggested that the real phenotype violated certain assumptions of the regression model. When we plotted the distribution of phenotype values, we saw that the real phenotype was more skewed than the BM phenotype and had longer tails that contained some extreme values (**Figure 1B**).

**Figure 1:**
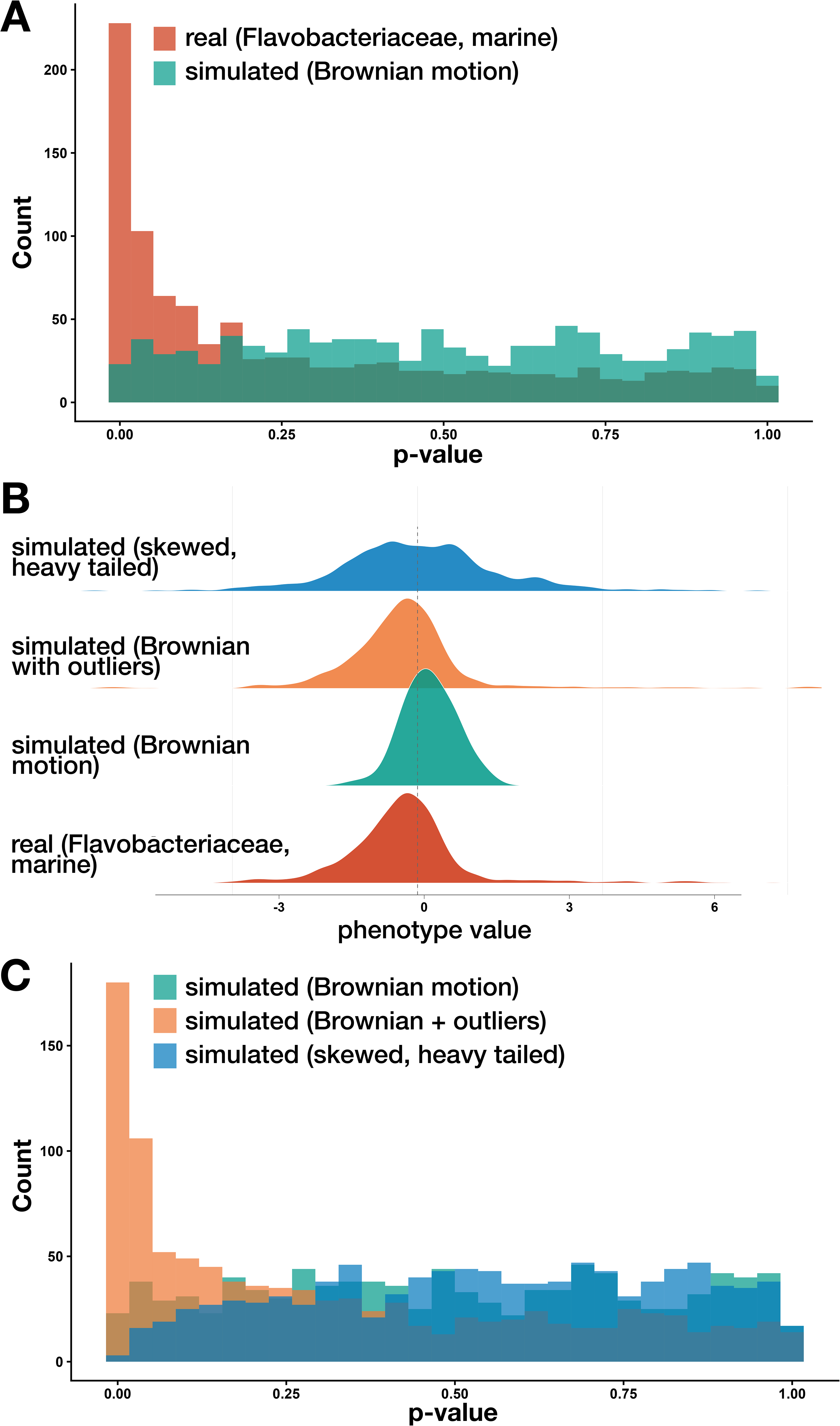
Phylogenetic linear regression is biased when phenotypes have outliers or violate distributional assumptions. A) Histogram of p-values under the null for a phenotype generated with Brownian motion (light) compared to a phenotype derived from an analysis of polar metagenomes (surface vs. mesopelagic zone, *Flavobacteriaceae*; dark). B) Density plots of simulated and real phenotypes. From top to bottom: Brownian motion rescaled to follow a skewed, heavy-tailed distribution (blue); Brownian motion with outliers added (orange); Brownian motion (tan); and the real phenotype from panel A (red). C) Histogram of p-values under the null for the three simulated phenotypes in panel B (same colors).

**Figure 2:**
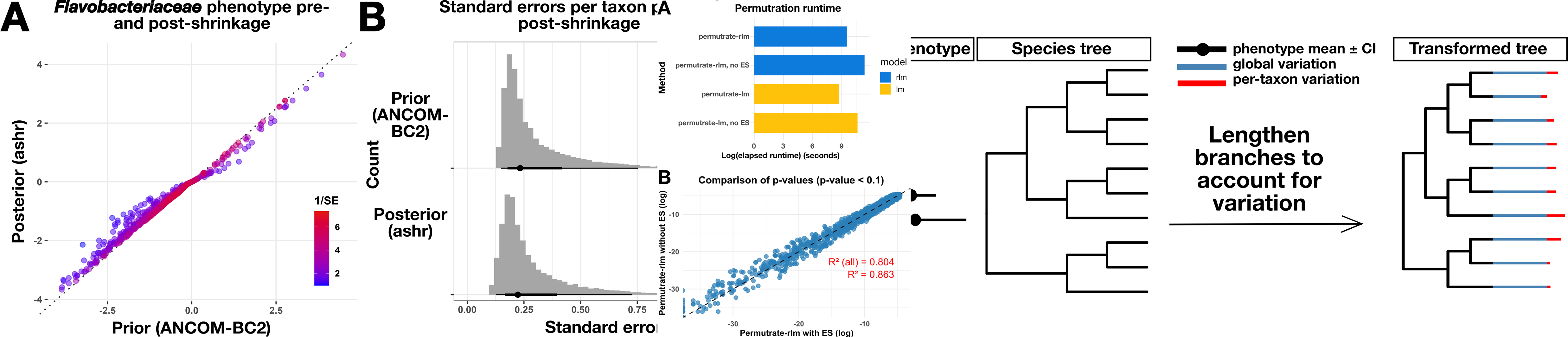
Integrating error via adaptive shrinkage and per-taxon branch extension. A) Scatter plot showing the differential abundance phenotype for *Flavobacteriaceae,* comparing mesopelagic abundance to surface abundance. Differential abundances were initially generated with ANCOM-BC2 (x-axis) and then shrunk using ashr (y-axis). Colors show the precision (inverse standard error) from red (more confidently measured) to blue (less confidently measured). B) Histograms and dot plots showing the distribution of standard errors from ANCOM-BC2 (top) and after applying adaptive shrinkage (bottom). C) Illustration of tree transformation that allows both global (blue) and per-taxon (red) variation to be integrated into a phylogenetic regression model. The right panel shows phenotype values that vary in both mean and standard error across the tree; the left shows the same tree, but with tips lengthened based on jointly fitting global variation and per-taxon variation (see Methods). Note that all tips are lengthened, but less confidently measured taxa are lengthened more.

To identify which of these could be leading to worse false positive control, we generated two additional phenotype sets: null BM phenotypes that were modified to introduce outliers, and null BM phenotypes that were rescaled to have elevated skew and kurtosis (**Figure 1B**; Methods). These simulations revealed two opposite failure modes of phylogenetic regression: anti-conservative behavior in the presence of phenotypes with outliers, and a conservative bias under high skew/kurtosis, indicating a potential loss of power (**Figure 1C**).

Together, these findings motivated the development of a new framework for *phylogenize* capable of handling microbial differential abundance data. Specifically, we wanted our improved method to have the following qualities:

1. Give accurate results even when outliers are present;
2. Account for measurement error;
3. Use an empirical null distribution instead of one based on, e.g., Brownian motion;
4. Remain computationally tractable (since it would be applied to hundreds of thousands of gene families).

### Development of “robust permutration”

#### Outlier tolerance

As we saw worse performance in the presence of outliers, we started by extending a recently published family of robust phylogenetic regression methods (ROBRT)^33^. ROBRT makes use of the fact that phylogenetic regression is equivalent to an older approach known as phylogenetic independent contrasts^41^ (PICs), in which the phylogenetic signal is removed from trait and predictor values before applying a standard linear regression. Because the resulting PICs are no longer correlated with the tree, this means they can also be analyzed using generic robust methods for regression, which use alternative estimators (e.g., M- and MM-estimators) that should be less influenced by outliers with high leverage. Robust phylogenetic methods have also been shown to be less sensitive to evolutionary model violations^33^ and tree misspecification^42^.

#### Integrating measurement error

Another possible shortcoming of phylogenetic regression is that most methods ignore the fact that different taxa may be measured with different precision. This is important for our application because our phenotypes are not measured directly, but rather propagated from a statistical metagenomic analysis. This partly motivated the development of shrinkage estimators for prevalence phenotypes in the original version of *phylogenize*^21,23^, which moved poorly measured estimates closer to zero (“no effect”), reducing their impact on the regression. Because of the additional statistical complications of abundance data, we decided to leverage the development of modern methods for analyzing microbiome data that report per-taxon standard errors^37,43^, which was not a feature of common earlier methods^44,45^. This allowed us to apply adaptive shrinkage^38^, a general-purpose empirical Bayes approach, instead of developing a novel shrunken estimator for abundances.

However, we observed that even after applying adaptive shrinkage, certain taxa remained much more confidently measured than others. This is difficult to account for with standard phylogenetic linear regression, which only allows for a single, global measurement error term. More flexible methods like phylogenetic generalized mixed models (PGLMMs^46^) can account for taxon-specific variation, but typically are more computationally intensive, requiring Markov Chain Monte Carlo algorithms to fit the model^25^. Integrating a robust estimator would also be much more complicated in a generalized mixed model framework than in the robust PIC framework noted above.

We therefore took an alternative approach based on tree transformation. Tree transformations, such as Pagel’s λ transformation^47^, use the fact that under the Brownian motion model, the variation in a trait between two nodes is proportional to the branch length connecting them. Non-phylogenetic measurement error can therefore be modeled without affecting the phylogenetic correlation structure simply by extending the branches at the tips of the tree. However, our approach differs from Pagel’s λ because we extend tips in proportion to two separate sources of error: one, like Pagel’s λ, is global, but the other is proportional to the standard error of each taxon (see Methods).

#### Generating an empirical, data-dependent null distribution

As noted, phylogenetic linear models typically use a neutral Brownian motion evolutionary model as a null distribution for traits. However, this may not capture (for example) variation in the evolutionary rate over time, “jumps” in trait values along a particular branch^32^, or traits with skewed or kurtotic distributions. An example of a method that uses an empirical null is “permulation,” which generates a null distribution with Brownian-motion simulations that are then rank-normalized to have the same distribution as the original trait^36^, thus combining <u>perm</u>utation and sim<u>ulation</u>. However, the use of a Brownian-motion simulation could potentially constrain the null distribution in the presence of jumps or variation in evolutionary rates over time. Inspired by the phylogenetic ridge regression^48^ method RRphylo^35^, we therefore also developed our own alternative empirical method called “permutration,” which comes from the <u>permu</u>tation of <u>rate</u>s on the tree. Like RRphylo, this method converts traits into per-branch rates using an ancestral state reconstruction; however, instead of directly performing an association test on these rates, we permute the rates and generate new null traits from them, allowing us to use the same PIC framework as above. To allow for variation in evolutionary rate over time, the branches are binned by their parent-node-to-root distance, and rates are only permuted within each bin (see Methods).

#### Computational tractability

Because these methods involve constructing an empirical null, which could be time-consuming when applied at scale, we implemented these approaches using an “early stopping” rule^49^. Briefly, this rule calculates the p-value continuously as more iterations are performed, then allows the test to terminate early if the p-value remains above a given (data-dependent) threshold. This rule is guaranteed under minimal assumptions to yield correct p-values under a user-specified threshold; by default, we set this to p=0.1, as it is unlikely that any p-values above a nominal threshold of 0.1 would be significant after correction. In addition to this rule, we also pre-compute null phenotypes and convert them to PICs, avoiding a potentially costly recalculation per gene family.

We find that despite allowing up to 1,000x more tests in the worst case, permutration is only around 20x slower than phylolm and 12x slower than POMS (**Figure 3A**), meaning that it remains tractable for large-scale analyses. Furthermore, the p-values with early stopping are highly correlated with those obtained without early stopping, especially (as expected) for values below the p=0.1 threshold (**Figure 3B**).

**Figure 3:**
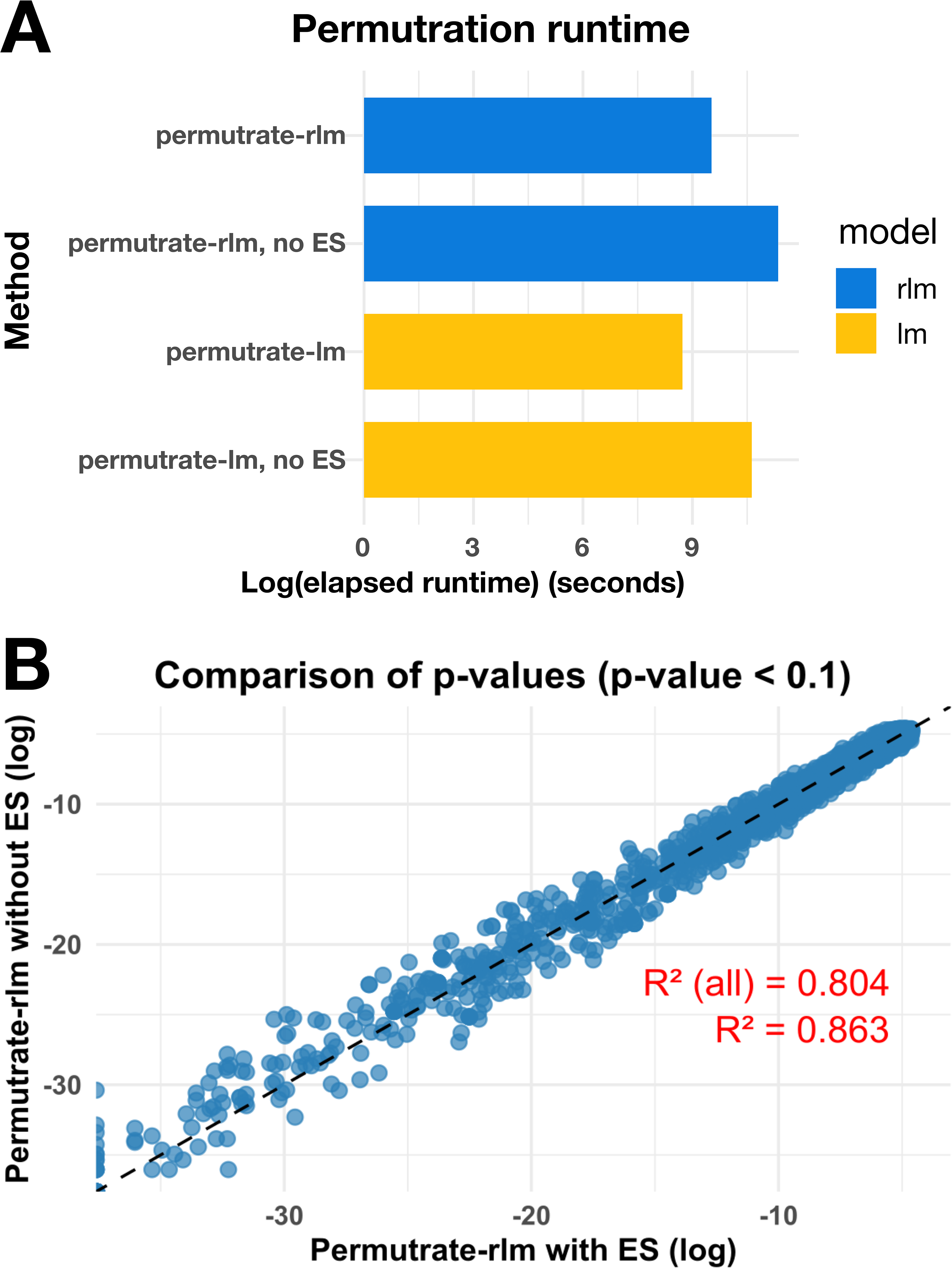
Early stopping makes permutration tractable at scale without affecting accuracy. A) Comparison of runtimes (x-axis) on simulated data for the permutration test, using either robust (rlm, blue) or standard (lm, yellow) linear regression, with or without early stopping (ES). B) Comparison of log p-values below the 0.1 threshold with (x-axis) and without (y-axis) early stopping. The dashed line shows equality. R^2^ values are reported for all p-values and for just those below the threshold, showing that the agreement is highest for sub-threshold p-values.

#### Overall evaluation

Combining the above changes yields a method we call “robust permutration.” We evaluated this method on synthetic metagenomic data, in which we simulated microbial abundances across two groups of samples. These simulations assume that 25 out of 1,000 genes truly contribute to changes in abundance. Briefly, we modeled the true abundances of each microbe using Brownian motion but then added asymmetry and heavier tails by rescaling this trait to have a skewed *t*-distribution; we then added additional global and per-taxon measurement noise, and in some cases, also added outliers (see Methods for details).

We compared true positive and false positive rates of permutration to phylolm (as used in the original *phylogenize*), ROBRT with the MM-estimator, and permulation, with and without the per-taxon tree transformation. We did not test ROBRT with the M-estimator because it does not yield p-values; however, for permutration and permulation, we were able to compare both robust (M-estimator) and non-robust (standard L2 linear regression estimator) versions, as the p-values are estimated empirically. Finally, we also compared to POMS, as well as a phylogenetic method developed for microbial GWAS called TreeWAS; TreeWAS implements three different metrics (“simultaneous”, “subsequent”, and “terminal”), all of which we tested.

We find that “robust permutration,” which uses robust linear modeling, tree rescaling, and permutrations, substantially improves control over the false positive rate compared to the standard phylogenetic linear model, while only incurring a small loss in power (**Figure 4**). Indeed, robust permutration was the most powerful method that appropriately controls the false positive rate (on average, 5% or lower at p = 0.05).

**Figure 4:**
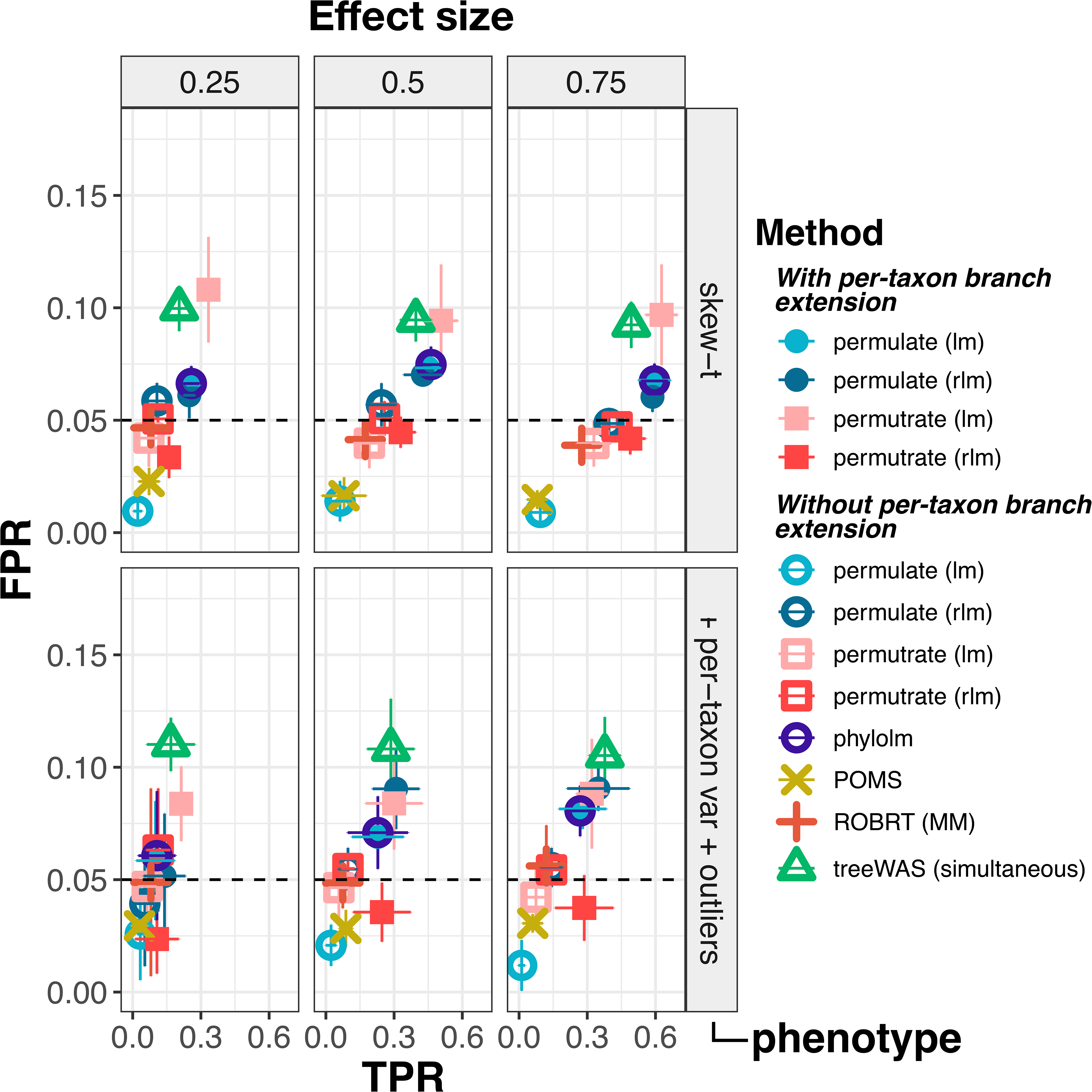
Robust permutration with per-taxon branch extension controls false positives while retaining more power than competing methods. The true positive rate (x-axis, TPR) and false positive rate (y-axis, FPR) are plotted for several methods applied to simulated datasets. The dashed line marks 5% false positives, the expected level at p = 0.05. In both rows, the true phenotype was simulated using Brownian motion and then quantile- normalized against a skew-t distribution (see Methods); in the bottom row, more per-taxon noise and five outlier points (out of 250; see Methods) were also added. Permulation and permutration were performed either with (filled shape) or without (open shape) per-taxon branch extension. In the legend, “rlm” refers to a robust linear model, “lm” to a standard linear model, and “phylolm” to phylogenetic regression. Full results including other TreeWAS methods and the uncorrected linear model are shown in Supplemental Figure 1 and Supplemental Figure 2.

Interestingly, even though permulation and permutration had performed similarly in our earlier tests, here, we saw a clear advantage for robust permutration under these more realistic simulations, especially in the presence of outliers and extra per-taxon variation.

In general, robust methods tended to have both higher power and lower false positives in these simulations, underscoring their value for analyzing metagenome-derived phenotypes. After robust permutration, the next best method that controlled the false positive rate was ROBRT-MM. However, ROBRT-MM also failed to converge frequently, resulting in many genes with no results, which was not an issue with the M-estimator methods. POMS had a low false positive rate, but also had low power, consistent with previous studies. Conversely, the “simultaneous” TreeWAS metric was competitive with robust permutration on power but consistently had among the highest false positive rates (the other two metrics both had low power; see Supplemental Figure 1). As expected, an uncorrected linear model had an extremely high false positive rate, at times reaching 40% (Supplemental Figure 2).

#### Case study: *Lachnospiraceae* that persist in the cirrhotic gut are enriched for an iron-sulfur transcription factor linked to oxygen detoxification

Several studies have found that liver cirrhosis is associated with a marked decline in gut commensals of the *Lachnospiraceae* family^1,50^, and that *Lachnospiraceae* abundance is anti-correlated with disease severity^1^. Furthermore, in mouse models of liver disease, including primary sclerosing cholangitis^51^ and alcohol-induced injury^52^, providing *Lachnospiraceae* strains has been observed to reduce disease severity. We were therefore interested in whether we could identify gene families that might play a protective role for *Lachnospiraceae* in the cirrhotic gut environment.

To do so, we re-analyzed gut metagenomes from a case-control study of cirrhosis (n=98 cases and 83 controls)^2^. A differential abundance analysis using ANCOM-BC2 revealed that while *Lachnospiraceae* tended to be depleted overall, others were much less so, with some even increasing in relative abundance. Applying robust permutration to these results, using the MGnify human gut database, revealed 242 significant positive associations at a q-value of 0.05.

In contrast, on the same dataset and database, a linear model found 9,391 significant positive associations, which is similar to the number of cirrhosis-associated orthogroups found in the original study (13,970). Protein families that were only significant in the linear model included highly conserved families like elongation factor Tu (linear *q* = 7.6 × 10^−13^, permutration *q* = 0.98) and DNA polymerase III (linear *q* = 3.0 × 10^−11^, permutration *q* = 0.34); this may be because certain clades that were more depleted had copies too divergent to be detected. They also clustered significantly more by clade than those only significant in Phylogenize2 or both models, indicating more potential for phylogenetic confounding (Ives-Garland α = 20 ± 0.24 vs. 39 ± 2.7 and 29 ± 1.8, respectively; p < 0.005). Furthermore, despite calling nearly 40-fold more gene families significant at q ≤ 0.05, the linear model still failed to recover 29% of the significant hits from robust permutration (70 out of 242). Finally, POMS found only 35 significant genes at q ≤ 0.05, only one of which (UniRef50_J9FR55) overlapped with robust permutration. Thus, robust permutration recovers almost seven times as many significant associations as POMS, but only 1/38th as many as the uncorrected linear model.

One of the most significant associations from the robust permutration analysis was a transcription factor in the Rrf2 family (UniRef50_C0FN14, q=0.0064). We also found weaker evidence of association for a second transcription factor in this family (UniRef50_A0A0J9BYM1, q=0.09; **Figure 5A**). Rrf2 transcription factors contain iron-sulfur (Fe-S) clusters, allowing many TFs in this family to sense oxidative and/or nitrosative stress. We would indeed expect reactive oxygen and nitrogen species to be increased under these conditions, as both intestinal permeability and systemic inflammation (potentially leading to the release of nitric oxide) tend to be higher in cirrhotic individuals^53^, and these factors have previously been linked to higher concentrations of oxygen, nitric oxide, and downstream metabolites^54^. These small molecules are known to have profound effects on the gut microbiome, creating niches for certain nitrate-respiring Proteobacteria while also inhibiting the growth of obligate anaerobes^54^ like the *Lachnospiraceae*.

**Figure 5:**
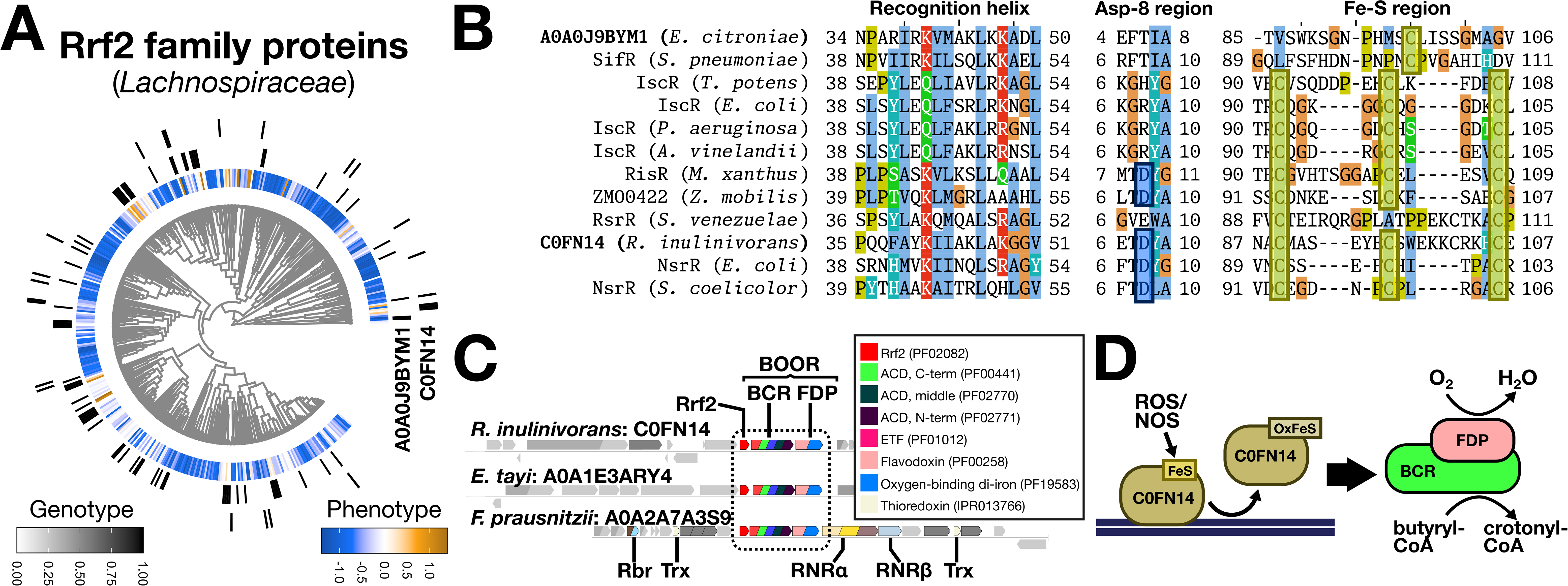
A transcription factor from *Lachnospiraceae* enriched in the cirrhotic gut is linked to oxygen detoxification. A) Phylogenetic tree of *Lachnospiraceae* showing log-fold-change in cirrhosis (red: higher abundance; blue: lower abundance) and the presence of two proteins in the Rrf2 family (A0A0J9BYM1, C0FN14). B) Multiple alignment of A0A0J9BYM1 and C0FN14 against known Rrf2 family sequences, showing in particular the recognition helix region, the presence or absence of an aspartate at position 8 (blue boxes), and the region where multiple conserved cysteine residues (yellow boxes) typically coordinate the Fe-S cluster. C) Gene neighborhoods of C0FN14 and two other homologs, A0A1E3ARY4 in *Eisenbergiella tayi* and *A0A2A7A3S9* in *Faecalibacterium prausnitzii.* A dashed box highlights the Rrf2 homolog and the butyryl-CoA-oxidoreductase (BOOR) system (comprising butyryl-CoA reductase, BCR, and the flavodiiron protein, FDP). Other domains and families of interest are colored (inset) and labeled (RNR: ribonucleotide reductase; Trx: thioredoxin; Rbr: rubrerythrin). D) Proposed function for C0FN14. We hypothesize that ROS/NOS oxidize the iron-sulfur cluster bound to C0FN14, causing it to detach from DNA and allowing BCR/FDP to be induced. This complex reduces molecular oxygen using electrons from butyryl-CoA.

We first constructed a multiple alignment against known Rrf2 sequences^55^ to determine whether C0FN14 and A0A0J9BYM1 likely coordinated Fe-S clusters (**Figure 5B**). These other Rrf2 sequences included IscR^56^, the main regulator of Fe-S cluster biosynthesis; RisR^57^, a diverged Rrf2 with a similar role to IscR; NsrR^58^, which typically senses NO (or, potentially in some organisms, nitrite); SifR, a quinone sensor^59^, and RsrR, a sensor of the general redox state. We confirmed that the three cysteine residues that typically coordinate the iron-sulfur cluster in Rrf2 proteins were conserved in C0FN14.

Additionally, C0FN14 had an aspartate at residue 8 that is known to be a fourth ligand for the Fe-S cluster in NsrR, and the recognition helix of C0FN14 resembled Rrf2 members like NsrR and RisR. In contrast, A0A0J9BYM1 appeared to only have a single conserved cysteine and more closely resembled SifR.

In aerobes, NsrR almost always regulates a flavohemoglobin protein that oxidizes NO to NO2; in *Streptomyces coelicolor*, this flavohemoglobin is in fact encoded by an ORF adjacent to NsrR. However, when we looked at the genome neighborhood of C0FN14, which is present in *Roseburia inulinivorans*, we instead found that it was encoded near a pair of genes annotated as an acyl-CoA dehydrogenase (ACD) and a flavodiiron protein (FDP). Further investigation using InterProScan^60^ revealed that this ACD had an unusual domain structure, with an ACD domain fused to an electron transfer flavoprotein (ETF) domain and a rubredoxin (RBR) domain. Interestingly, a protein with this exact domain structure, called BCR (for butyryl-CoA reductase), has previously been studied in the organism *Fusobacterium nucleatum*, where it is also encoded next to an FDP. That study showed that BCR and FDP together function as a butyryl-CoA oxygen oxidoreductase (BOOR), coupling the oxidation of butyryl-CoA to crotonyl-CoA with reduction of molecular oxygen^61^. Furthermore, the authors also showed that outside of *Fusobacterium*, the BOOR gene pair was conserved in anaerobic, butyrogenic gut Firmicutes, including *R. inulinivorans*, giving us confidence that this represents a legitimate BOOR system.

We generated a sequence-similarity network for C0FN14 using the Enzyme Function Initiative (EFI) web tools^62,63^, which showed that close homologs were overwhelmingly found in gut Clostridia, especially the families *Oscillospiraceae* (in particular, *Faecalibacterium prausnitzii*), *Acutalibacteraceae*, *Eubacteriaceae*, and *Lachnospiraceae*. Using the EFI genome neighborhood tool, we then looked for combinations of Pfam domains^64^ that were frequently represented in neighboring genes. This revealed that close homologs of C0FN14 were almost all encoded adjacent to BOOR gene pairs. Other proteins enriched in these neighborhoods, especially in *F. prausnitzii*, also included rubrerythrin, thioredoxin, and both subunits of ribonucleotide reductase (RNR; **Figure 5C**). Rubrerythrin is a peroxide scavenger unrelated to catalase and superoxide dismutase, which has previously been shown to provide oxidative stress resistance in the Gram-negative *Porphyromonas gingivalis*^65^.

Thioredoxin is a small redox protein that often acts as the electron donor for RNR systems^66^. Interestingly, aerobic class I RNRs have also been shown to improve oxygen resistance in the Gram-negative anaerobe *Bacteroides fragilis*^67^, and have also been observed in the *Fusobacterium* genome^68^.

These results point to a role for C0FN14 in regulating the oxidative stress response in more resistant *Lachnospiraceae,* as well as potentially in *Oscillospiraceae* like *F. prausnitzii*. In Gram-negative gut taxa like the Bacteroidia, transcriptional regulators of the OxyR and PerR families have been shown to control the response to oxygen, with some evidence of horizontal transfer and even of recent selection for increased oxygen tolerance^69^. However, OxyR and PerR are not members of the Rrf2 family, and OxyR in particular does not appear to be conserved in the *Lachnospiraceae*; C0FN14 therefore appears to represent a novel component of this response in gut Clostridia. These results are also informative about the environment: they indicate that oxidative stress is likely a major selective pressure for microbes in the cirrhotic gut, and point to the BOOR system, RNR, and rubrerythrin as mechanisms by which anaerobic gut Clostridia may better withstand this stress. We finally note that, despite being one of the most significant associations with cirrhosis found by robust permutration (q=0.0064), this transcription factor was not identified by POMS (q=0.81) and was ranked in the bottom half of the 9,391 associations from the linear model (q=0.012).

## Conclusions

In realistic simulations of metagenomic abundance data, we found that robust permutration was the most sensitive phylogenetic comparative approach that retained good control over the false positive rate. Robust permutration was developed to address the specific challenges of these data, such as the presence of outliers and taxon-specific measurement error. We also found that on real data, robust permutration recovered substantially more gene associations than the only other phylogenetic method for microbiome data, POMS. At the same time, it remained much more selective than a standard linear model, which tended to return thousands of additional genes with high phylogenetic confounding. We have implemented robust permutration in an open-source R package (https://github.com/pbradleylab/repermulize) and have also made it the default method in a revised software package, Phylogenize2, allowing it to be applied across diverse environments (see Companion).

POMS uses the phylogenetic isometric log-ratio transform (PhILR^34^) to deal with the compositional nature of microbiome data; this transforms the per-taxon abundances to “balances” at each node in the tree. Conceptually, these are very similar to the independent contrasts used in PIC, raising the question of why we see lower sensitivity with POMS. We believe that two main factors are responsible. First, as the authors of POMS noted, POMS uses rank-based tests to compare branches from each node; nodes near the tips of the tree will therefore have lower sample sizes, lowering sensitivity for recent changes. Second, POMS requires a statistically significant association between the output of two rank-based tests, which are individually binarized at a nominal p=0.05. This likely results in a conservative bias. It may be possible to use an approach like permutration to empirically recalibrate the results of POMS, as well as some of the other comparative methods we tested, such as the non-parametric statistics derived from TreeWAS.

We have also shown that the results from applying robust permutration to real data can point us to specific hypotheses that are biologically coherent, such as the putative oxidative stress regulon we identified in *Lachnospiraceae*. Comparative methods capable of identifying individual genes of interest may be especially valuable in clades that are currently (for the most part^70^) genetically intractable, yet have strong links to health, like the *Lachnospiraceae*^51,52,71,72^. These results also suggest that certain commensal microbes may be better suited to colonize the gut in cirrhosis or high-fat diets because of their intrinsic resistance to oxidative and/or nitrosative stress. On the other hand, we would expect the BOOR system to also reduce butyrate production, since butyryl-CoA is the substrate. Butyrate is both an energy source and signaling molecule in the gut epithelia and promotes improved barrier integrity, combating the effects of inflammation^54,73^. Thus, even though *Lachnospiraceae* are in general associated with better liver health^1^, and some do appear to survive better in the cirrhotic gut, not all of these may be equally effective in mitigating disease symptoms. Identifying more stress-resistant microbes that are still capable of producing key metabolites such as butyrate could potentially help prioritize strains for investigation as new probiotics.

## Methods

### Metagenomic data processing

Three shotgun metagenomic datasets representing distinct biomes were retrieved from the European Nucleotide Archive^74,75^. These were: 1) a case-control study of the human gut microbiome in cirrhosis (accession PRJEB6337)^2^; 2) an experimental study of the mouse gut microbiome on normal vs. high-fat diets (PRJEB52043)^76^; and 3) polar marine microbiome samples from the Tara Oceans project (PRJEB9740), focusing on the prokaryotic size fraction at two ocean depth strata, the surface (SRF) and mesopelagic (MES) zones.

All sequencing reads underwent quality control and adapter trimming using FastQC and MultiQC^77^ (both with default settings), and BBDuk (right-quality trimming at a Phred score threshold of 10). Taxonomic profiling was performed using Kraken 2^78^ using the human gut, mouse gut, and marine MGnify reference databases, respectively; species counts were then estimated from these profiles with Bracken^79^. For the human samples, some subjects had multiple sequencing runs, which were merged via summation prior to analysis. We used ANCOM-BC2^37^ to estimate differential abundances and standard errors for each taxon, comparing (respectively) cirrhosis controls to cases, mice on normal chow to those on high-fat chow, and marine mesopelagic to surface samples. Finally, adaptive shrinkage (using the R package ashr^38^) was used to shrink these estimates, yielding a posterior effect size and standard error. These were used as the per-taxa “phenotype” for downstream analyses.

### Initial test of phylogenetic linear model

To assess the impact of modeling assumptions and outliers on the phylogenetic linear model, we simulated 72,545 gene values that were uncorrelated with our phenotype of interest, meaning that the null hypothesis of no association should be true. Continuous null genes were simulated along the phylogenetic tree using the Brownian motion model implemented in the phylolm R package^30^, generating continuous gene trait values. We then used phylolm to associate these with either simulated phenotypes or the values for *Flavobacteriaceae* from the marine dataset. In both cases, we used the tree from the MGnify marine collection, made ultrametric using relative evolutionary divergence^80^ as implemented in the castor R package^81^. Simulations were performed under the following scenarios:

1. Brownian motion, using the rTraitCont function from ape R package, the sigma parameter set at 1, and the model of “BM”, as implemented in phylolm.
2. Brownian motion with outliers. Simulated phenotypes were perturbed by replacing 10 randomly selected values with extreme values exceeding four standard deviations from the mean (z-score > 4).
3. Brownian motion, followed by quantile-normalization against a skewed t-distribution using the rskt function (and gamma_skew parameter at 2) from the skewt package, to introduce additional skew and kurtosis.

Finally, phylogenetic linear modeling as implemented in phylolm was performed to associate genes and phenotypes, and the resulting p-value distributions were compared (**Figure 1**).

“Robust permutration” pipeline

### Incorporating taxa-specific uncertainty into species tree

Due to measurement noise and sampling variability, taxa differ in how accurately their phenotypes are being estimated. Therefore, we model this uncertainty by estimating taxon-specific standard deviations (SDs) and incorporating them into the phylogeny (Fig. 2 – step 1). We first standardize *s_i_* , the SD for taxa *i*, as:

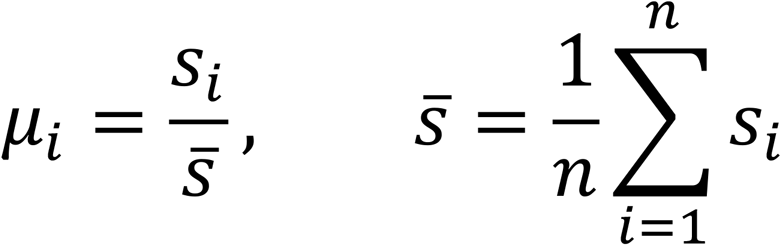

We next define a mixing parameter *m* controlling the balance between shared non-phylogenetic variation and taxon-specific uncertainty, as shown below, where *μ*^∗^ is the total uncertainty for taxon *i*, and *m_L_* is the logistic transform of *m*:

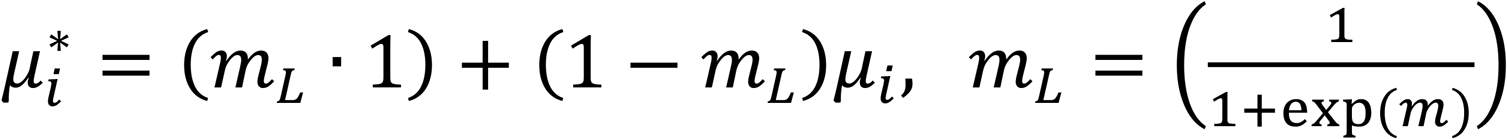

When *m_L_* = 0, the additional variation is purely driven by the taxon-specific variation, while at *m_L_* = 1, we ignore measurement heterogeneity completely, leading to a model similar to Pagel’s *λ*. Finally, we also define a global scaling factor *α*, representing how much to scale the total uncertainty:

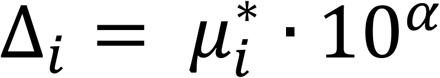

The parameters *α* and *m* are fit using Nelder-Mead optimization (implemented in R) to maximize AIC of a Brownian motion model fit to the phenotype data (using phylolm). The resulting per-taxon uncertainty values are then used to proportionally extend branch lengths:

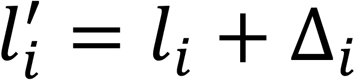

where *l_i_* is the original branch length and *l*^2^ is the extended branch length. Conceptually, after lengthening, taxa with less precise estimates or higher uncertainty, will have longer branch as they evolve more independently, and thus contribute less to the overall phylogenetic structure.

### Estimating an empirical null with permuted evolutionary rates

Traditional permutations erase phylogenetic structure^36^. One approach that avoids this problem is permulation^36^, which simulates data under Brownian motion and then rank- normalizes to the original data. As an alternative, inspired by approaches like RRphylo^35^, we instead convert trait values to rates of change at each branch of the tree. These rates, like PICs, should be independent of phylogenetic structure. Unlike RRphylo, though, we do not directly fit a model on these rates. Instead, we permute them and then use the permuted rates to generate new random traits, which allows us to construct an empirical null distribution.

To obtain rates, we first perform ancestral state reconstruction of the phenotype on the rescaled phylogenetic tree (see above) using the phylogenetic independent contrasts (PIC) framework^41^, implemented in the *castor* R package^81^. This estimates trait values for internal nodes under a Brownian motion model of evolution. We then convert evolutionary changes along each branch *i* into standardized evolutionary rates *r_i_* by computing the difference in trait values Δ*x_i_* between connected nodes and scaling them by the square root of the branch length, 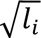:

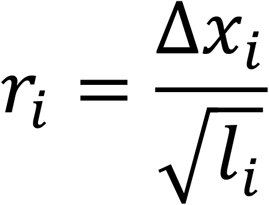

Branches are then grouped into bins (by default, 10 bins) based on the node-to-root distance of the parent node. Rates are then permuted within each bin, allowing for changes in the distribution of evolutionary rates over time. Finally, these rates are rescaled by multiplying by the root branch length, yielding new trait values 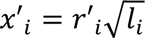.

Empirical p-values are calculated by first converting real trait and genotype values to PICs. Then, as in ROBRT^33^, we calculate the effect size of association between trait and genotype PICs using the robust M-estimator, as implemented in the rlm function from the MASS R package^82^. Next, an empirical null distribution of effect sizes is constructed by performing the same association with the permulated traits (by default, 1,000 permulations).

We implement an early-stopping rule^83^ to control the runtime, which estimates a running p-value and compares it to a cutoff *k*:

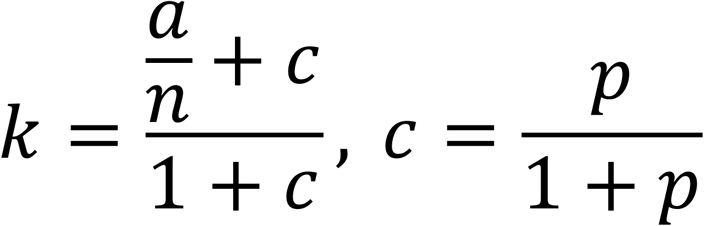

Over successive permutations *n*, *k* asymptotically approaches a fixed value *p* at a rate governed by the magnitude of *a*. By default, we set *p* to 0.1 and *a* to 10. This yields p-values that are accurate below 0.1, and guarantees at least 10 permutations will be tested before there is any chance of early stopping.

To evaluate how efficiently early stopping optimizes runtime (**Figure 3**), we used phenotype values from the *Flavobacteriaceae* marine dataset and simulated 72,525 null gene values. Phylogenize2’s permutation procedure (with rlm or lm) was run on this genotype-phenotype set with either the default early stopping parameter *a* = 10 or with *a* = 2,000 (which disables early stopping, as the cutoff will remain above a p-value of 1 for all 1,000 iterations). Runtime for each run was recorded and the elapsed runtimes and p-values were compared across settings.

Finally, because true associations are typically very far from the null, and thus could require a very large number of permutations to compute exactly, we obtain our final p-values using a normal approximation of the null: that is, we perform a z-test of the real trait value vs. the null trait distribution. We also tested a non-parametric approach using a kernel density estimate of the null distribution but found that it yielded similar p-values.

A parallelized version of this test is implemented in the package “repermulize,” which we have made available on GitHub (https://github.com/pbradleylab/repermulize). This package also includes our own implementation of permulation.

### Comparisons against other methods

We performed a systematic comparison of a large number of phylogenetic and non-phylogenetic approaches on simulated metagenomic abundance data. Simulations were performed using the following algorithm:

1. A random coalescent tree with 250 species is generated using the rcoal function in the ape R package.
2. Presence/absence profiles for 1,000 genes are simulated:

a. First, 5,000 profiles are simulated using the rbinTrait function in phylolm^30^. The *α* and *β* parameters were drawn randomly from the *U*(0.1, 30) and *U*(−2, 2) uniform distributions, respectively.
b. Next, these profiles are filtered to exclude genes with fewer than 3 presence values and fewer than 3 absence values.
c. Finally, 1,000 of the remaining profiles are randomly sampled. Out of these, 25 are selected as “target” genes (meaning they confer an effect on differential abundance of taxa).
3. Taxon abundances are simulated:

a. The “baseline” log-abundance for each taxon is drawn randomly from a *t-* distribution centered on 0 with 5 degrees of freedom.
b. Differential abundance is simulated as a combination of true effects and residual variation:

i. Residual phylogenetic variation is initially simulated with Brownian motion using the rtrait function from phylolm^30^, using an ancestral state of 0 and a variance of 1. Next, it is rescaled using rank-normalization to have a skewed t-distribution with 10 degrees of freedom and skewing parameter *γ* = 2, which is generated using the skewt R package^84^.
ii. To model true effects, for each of the 25 “target” genes, any taxon containing that gene has a constant added to its differential abundance (given by the effect size parameter).
c. Abundance vectors are converted to counts in 100 samples with 50 per group using a multinomial-Dirichlet approach, optionally adding extra non-phylogenetic variation per sample:

i. The total number of reads per sample is drawn from a lognormal distribution (log mean = 16.5, log SD = 0.5).
ii. In the first group, the baseline abundance vector is used, while in the second group, the baseline abundance vector plus the differential abundance vector is used.
iii. Extra non-phylogenetic noise is added to the abundance vector for each sample, centered on 0; the standard deviation is an adjustable parameter.
iv. Abundance vectors are converted to relative abundances.
v. For each sample, a multinomial draw from the relative abundance vector is taken (using the number of reads for that sample). A Dirichlet sample is then taken from this multinomial draw and scaled to the number of reads for that sample.
4. Differential abundance values are calculated from the count matrix.

a. Either MaAsLin2^43^ or ANCOM-BC2^37^ is used to compare groups, returning effect sizes and standard errors.
b. Posterior effect sizes and standard errors are computed using adaptive shrinkage (ashr)^38^, yielding a phenotype vector.
c. If desired, outliers are added to the phenotype:

i. Data points at least 1 standard deviation from the mean phenotype value are randomly selected to convert to outliers.
ii. Selected points are moved an additional 2 standard deviations away from the mean phenotype value (higher for data points above the mean, lower for data points below the mean).
5. Each phylogenetic method is applied to the same simulated dataset and evaluated for false positive rate (fraction of non-target genes with p≤0.05) and power (fraction of target genes with p≤0.05).

Each of 18 methods was evaluated in 3 separate sets of simulated trees and genes, with 3 random phenotypes generated per tree/gene set, at each of 24 parameter combinations. The parameters we varied were the differential abundance method (ANCOM-BC2 or MaAsLin2), extra non-phylogenetic variation (from a log-normal distribution with either *μ* = −10, *σ* = 2, effectively adding no extra variation, or *μ* = −2, *σ* = 2), the effect size (0.25, 0.5, or 0.75), and the number of outliers (0 or 5). However, certain simulations at some parameter combinations failed and were excluded, for example, because fewer than 3 taxa were found to be differentially abundant, because ashr could not fit the results (usually indicating not enough variation in the trait), or because the method returned another error. This was most common at the lowest effect sizes, where signal was weakest. The final dataset therefore consisted of 3,451 simulated runs; each method had at least 181 runs except POMS, which had 116.

The methods we tested comprised:

1. Linear regression;
2. The phylogenetic linear model (phylolm);
3. POMS;
4. TreeWAS, using the “simultaneous”, “subsequent”, and “terminal” statistics and either the original or rank-transformed data (6 combinations);
5. ROBRT with the MM-estimator;
6. PIC with permulation and early stopping (with or without the M-estimator); and
7. PIC with permutration and early stopping (with or without the M-estimator).

Finally, all methods were tested either using the original tree (allowing for measurement error either through the method itself, as with phylolm, or by using Pagel’s *λ* transformation^47^ as implemented in phylolm^30^), or the tree with branches lengthened to model per-taxon variation (see above).

As noted, we did not test ROBRT with the M-estimator because it does not return p-values; we also did not test the permulation/permutration methods with the MM-estimator because this estimator required significantly more compute time.

In **Figure 4**, we chose to present only the non-rank-transformed results from TreeWAS, since the results were similar to the rank-transformed results; we also only showed the results from the “simultaneous” metric as the others both had very low power on our simulated traits. We also did not present the results from the linear model as the false positive rates were very high (approaching 40%) and were difficult to visualize on the same scale. Finally, we presented only the results from ANCOM-BC2 in **Figure 4**, as we found a slight anti-conservative bias with the use of MaAsLin2. The full results are included in Supplemental Figure 1 and Supplemental Figure 2.

### Human gut in cirrhosis: analysis of Rrf2 family proteins

We performed the robust permutration test on abundances of *Lachnospiraceae* in cirrhosis (estimated as above). This test also requires estimates of protein family content and a phylogenetic tree, which were derived from the MGnify Biomes “human gut” database integrated into the latest release of Phylogenize2 (for details, see companion paper).

A multiple alignment of representative Rrf2 sequences was constructed using MUSCLE v5 via the EMBL-EBI Job Dispatcher^85,86^ and visualized in JalView^87^.

A sequence-similarity network for C0FN14 was generated through the Enzyme Function Initiative (EFI) online tool^63^, using an initial sequence similarity score cutoff of 10, resulting in a fully connected network of 202 sequences. Cutoffs were then adjusted interactively using Cytoscape^88^ to a final cutoff of 50, yielding 9 clusters; C0FN14 was a member of the largest, cluster 1 (151 sequences). Using the EFI genome neighborhood network (GNN) tool, we identified the most common Pfam domain architectures in proteins encoded near these cluster 1 sequences (149 neighborhoods). The most common domain architecture was “none,” indicating no annotations (in 123 neighborhoods), followed by “Rubredoxin-Acyl-CoA_dh_1-ETF_alpha-Acyl-CoA_dh_M- Acyl-CoA_dh_N” (the BOOR BCR, 123 neighborhoods) and “Flavodoxin_1-ODP” (the BOOR FDP, 115 neighborhoods). The full table can be found in Supplementary Table 2.

## Availability

The robust permutration test is available through https://github.com/pbradleylab/repermulize and is implemented in Phylogenize2 (https://github.com/pbradleylab/phylogenize; see Companion). Scripts for figure generation are available at https://github.com/pbradleylab/phylogenize_v2_2025_paper. Metagenomic simulation code is available at https://github.com/pbradleylab/phy2tests.

## Acknowledgements

We wish to acknowledge Dr. Kirsten Wolthers for providing insight on the function and distribution of the BOOR system, as well as other members of the Bradley lab for helpful discussions. Funding for this study was provided by The Ohio State University (startup funds to P.H.B.) and by the National Institutes of Health (grant R35GM151155 to PHB). The Ohio Supercomputer Center provided high-performance compute resources.

## Supplemental Figure and Table Legends

**Supplemental Figure 1:** **Full results of simulations from Figure 4 (excluding uncorrected linear regression).**

**Supplemental Figure 2:** **Full results of simulations from Figure 4 (including uncorrected linear regression).**

**Supplemental Figure 3:** **Two top hits specific to the uncorrected linear model applied to Lachnospiraceae in liver cirrhosis, EF-Tu and DNA polymerase III.**

**Supplementary Table 1:** Positive gene associations with human gut *Lachnospiraceae* in cirrhosis that were significant (q<0.05) via robust permutration (“perm”), the linear model (“lin”), and/or POMS (“poms”).

**Supplementary Table 2:** Genome neighborhood network analysis for C0FN14 showing overrepresented Pfam domain architectures. Domain architectures that match a protein discussed in the paper are noted in the “annotation” column.

