## Supplementary figures and images for "A robust, sensitive phylogenetic method enables gene-level metagenomic analyses"

### Supplemental Figure 1

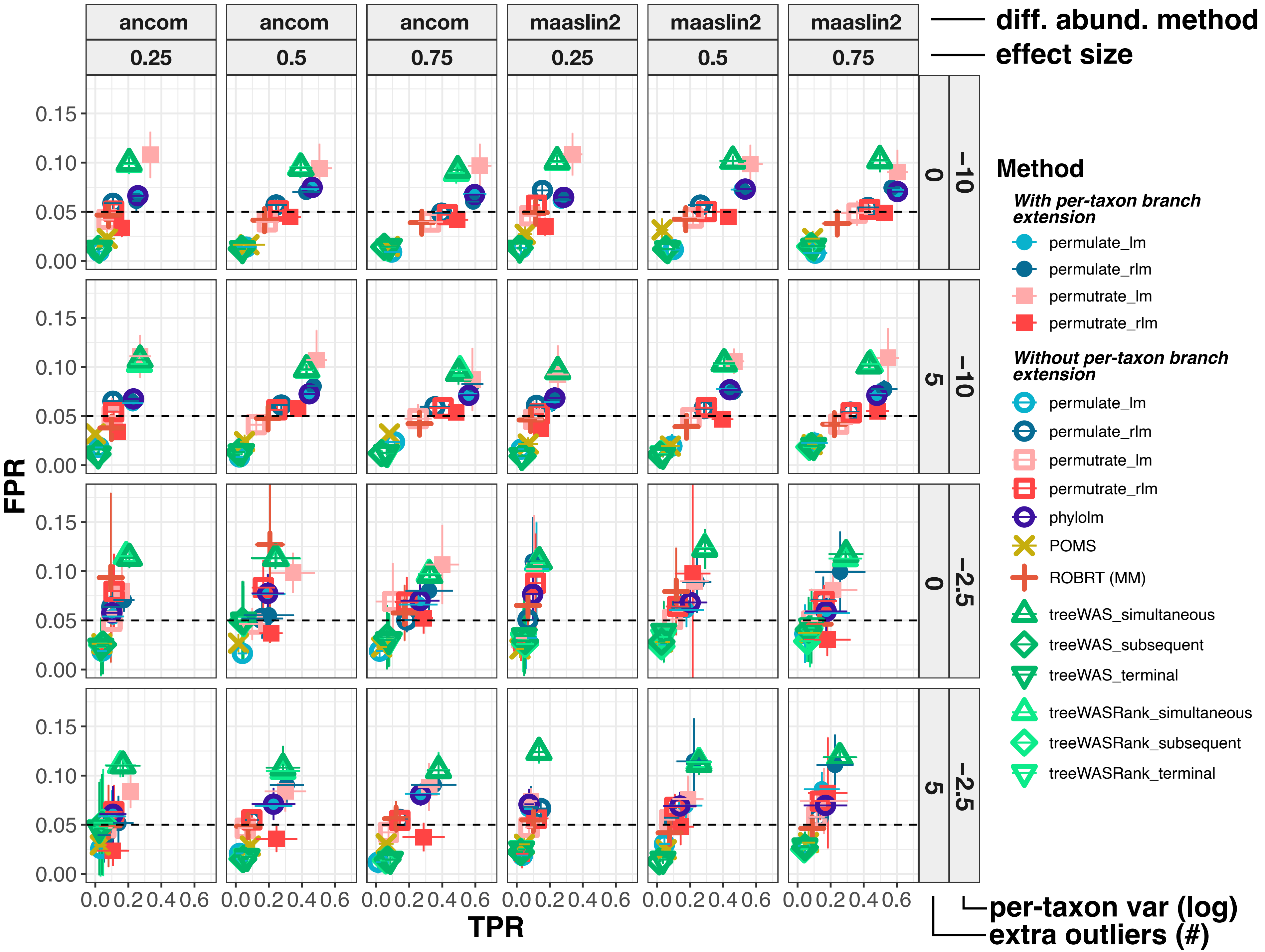

### Supplemental Figure 2

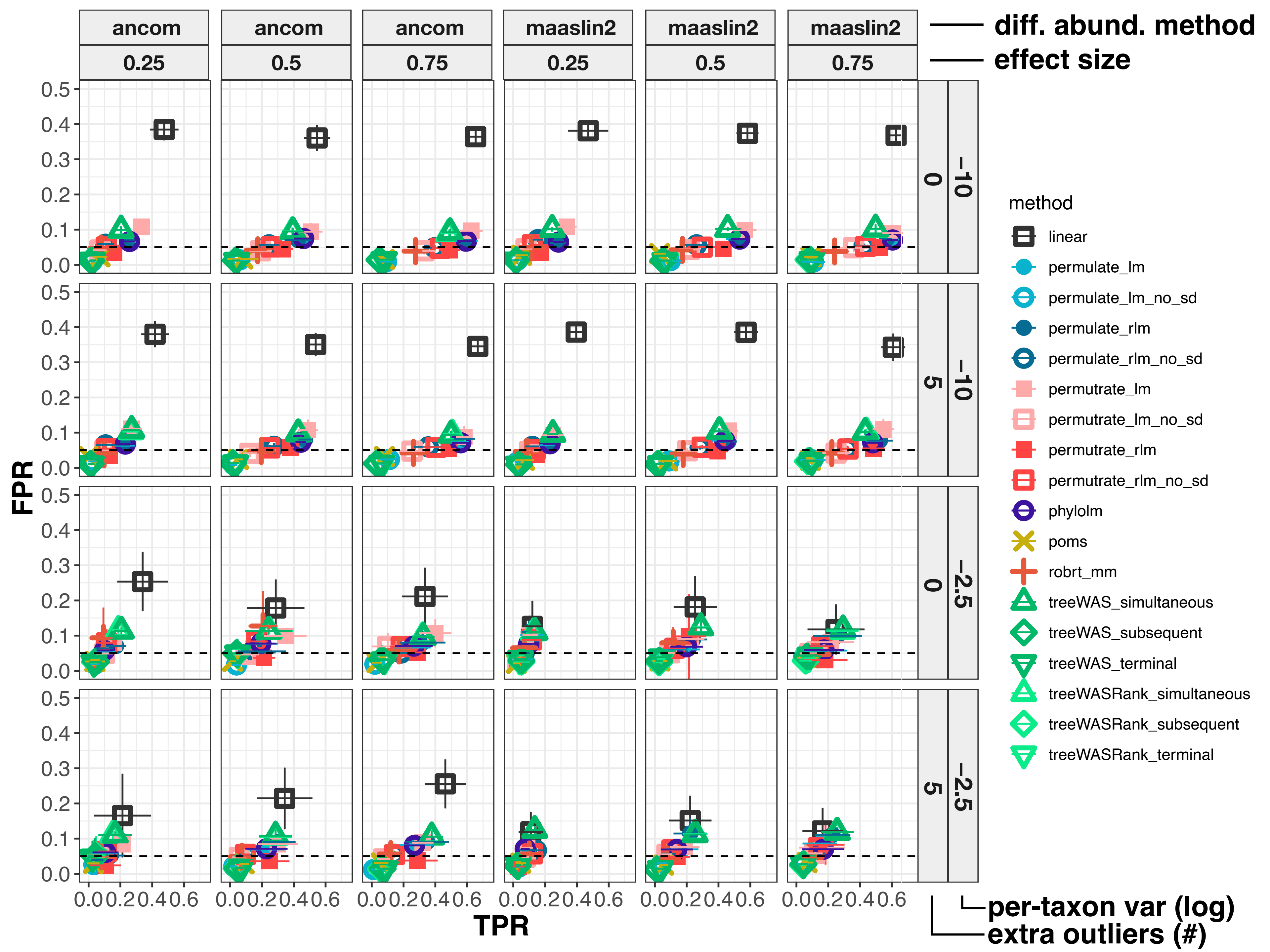

### Supplemental Figure 3

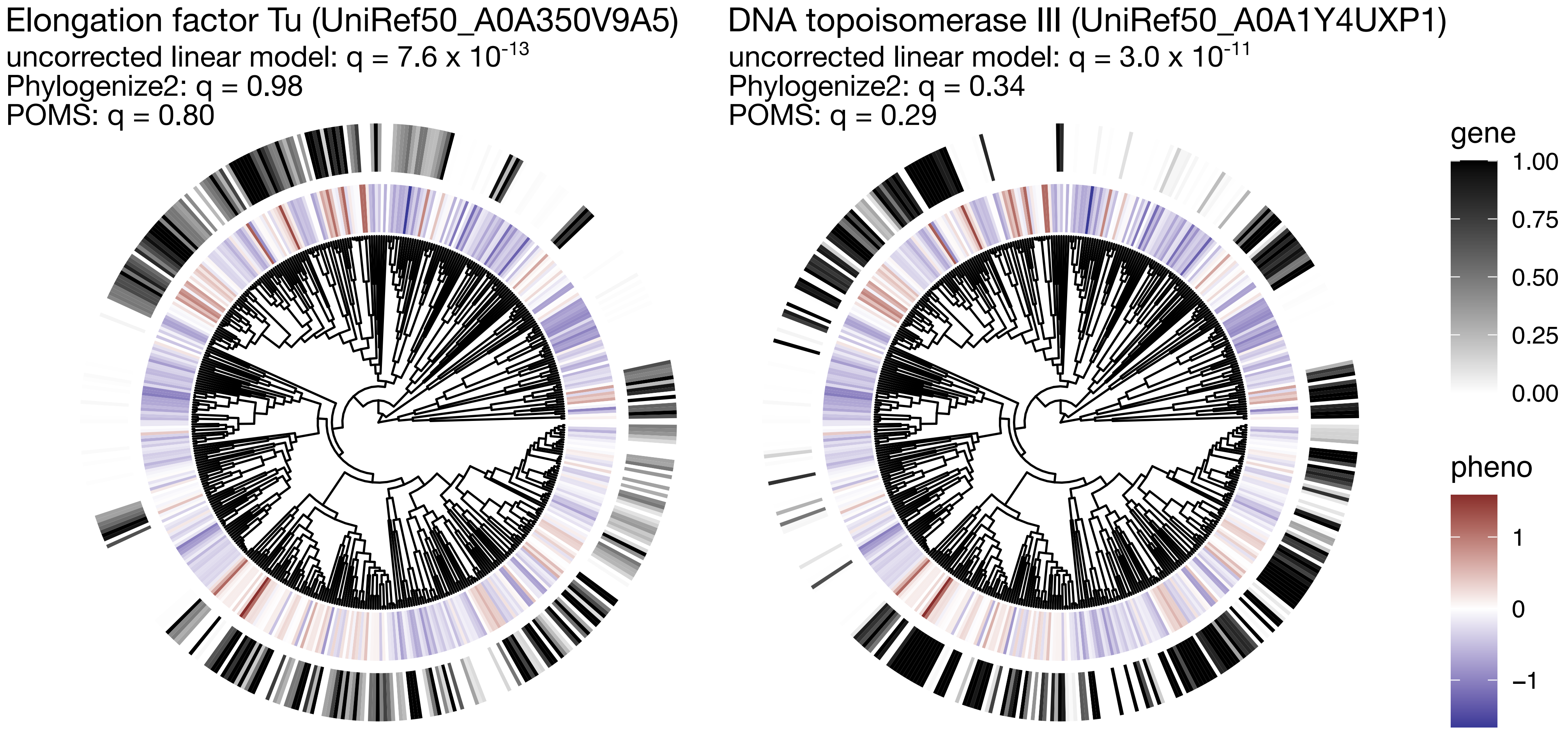
